# Synthetic metabolic pathways for photobiological conversion of CO_2_ into hydrocarbon fuel

**DOI:** 10.1101/381913

**Authors:** Ian Sofian Yunus, Julian Wichmann, Robin Wördenweber, Kyle J. Lauersen, Olaf Kruse, Patrik R. Jones

## Abstract

Liquid fuels sourced from fossil sources are the dominant energy form for mobile transport today. The consumption of fossil fuels is still increasing, resulting in a continued search for more sustainable methods to renew our supply of liquid fuel. Photosynthetic microorganisms naturally accumulate hydrocarbons that could serve as a replacement for fossil fuel, however productivities remain low. We report successful introduction of five synthetic metabolic pathways in two green cell factories, prokaryotic cyanobacteria and eukaryotic algae. Heterologous thioesterase expression enabled high-yield conversion of native acyl-ACP into free fatty acids (FFA) in *Synechocystis sp*. PCC 6803 but not in *Chlamydomonas reinhardtii* where the polar lipid fraction instead was enhanced. Despite no increase in measurable FFA in *Chlamydomonas*, genetic recoding and over-production of the native fatty acid photodecarboxylase (FAP) resulted in increased accumulation of 7-heptadecene. Implementation of a carboxylic acid reductase (CAR) and aldehyde deformylating oxygenase (ADO) dependent synthetic pathway in *Synechocystis* resulted in the accumulation of fatty alcohols and a decrease in the native saturated alkanes. In contrast, the replacement of CAR and ADO with *Pseudomonas mendocina* UndB (so named as it is responsible for 1-undecene biosynthesis in *Pseudomonas*) or *Chlorella variabilis* FAP resulted in high-yield conversion of thioesterase-liberated FFAs into corresponding alkenes and alkanes, respectively. At best, the engineering resulted in an increase in hydrocarbon accumulation of 8- (from 1 to 8.5 mg/g dell dry weight) and 19-fold (from 4 to 77 mg/g cell dry weight) for *Chlamydomonas* and *Synechocystis*, respectively. In conclusion, reconstitution of the eukaryotic algae pathway in the prokaryotic cyanobacteria host generated the most effective system, highlighting opportunities for mix-and-match synthetic metabolism. These studies describe functioning synthetic metabolic pathways for hydrocarbon fuel synthesis in photosynthetic microorganisms for the first time, moving us closer to the commercial implementation of photobiocatalytic systems that directly convert CO_2_ into infrastructure-compatible fuels.

**Highlights:**

- Synthetic metabolic pathways for hydrocarbon fuels were engineered in algae
- Free fatty acids were effectively converted into alkenes and alkanes
- Transfer of algal pathway into cyanobacteria was the most effective
- Alkane yield was enhanced 19-fold in *Synechocystis spp*. PCC 6803
- Alkene yield was enhanced 8-fold in *Chlamydomonas reinhardtii*

## INTRODUCTION

Cyanobacteria (prokaryotes) and algae (eukaryotes) are photosynthetic microorganisms that have evolved to naturally accumulate C15-C19 alkanes or alkenes at very low concentrations (0.02-1.12% alkane g/g cell dry weight (cdw)) (Lea-Smith et al., 2015; Schirmer et al., 2010; Sorigué et al., 2017) with the exception of naturally oleagineous species (Ajjawi et al., 2017; Metzger and Largeau, 2005; Peramuna et al., 2015). These hydrocarbons are postulated to influence the fluidity of cell membranes and are therefore essential for achieving optimal growth, indeed, the abolition of their biosynthetic capacities results in morphological defects (Lea-Smith et al., 2016). Only two enzymes, acyl-ACP reductase (AAR) and aldehyde deformylating oxygenase (ADO) are required to catalyze the bacterial conversion of acyl-ACP into alkanes (Schirmer et al., 2010). Similarly, eukaryotic microalgae also biosynthesize small quantities of alkanes and alkenes directly from fatty acids, employing the distinctly different and recently discovered fatty acid photodecarboxylase (FAP; (Sorigué et al., 2017)).

In order to engineer sustainable biotechnological systems for production of hydrocarbons for the fuel market, whether heterotrophic or light-driven, far greater yields are needed alongside other complementary non-biochemical improvements such as improved bio-process designs. Several studies have attempted to enhance alkane productivity in cyanobacteria by over-expression of the native or non-native AAR and ADO enzyme couple (Hu et al., 2013; Kageyama et al., 2015; Peramuna et al., 2015; Wang et al., 2013) which relies on acyl-ACP as the precusor. Although naturally accumulating alkane amounts have been enhanced through engineering and reported in high titres from the lipid-accumulating cyanobacteria, *Nostoc punctiforme* (up to 12.9% (g/g) cdw, (Peramuna et al., 2015)), similar efforts in the non-lipid accumulating model cyanobacterium *Synechocystis sp*. PCC 6803 (hereafter *Synechocystis* 6803) have at best yielded only 1.1% (g/g) cdw (Hu et al., 2013; Wang et al., 2013). In eukaryotic algae, the native alkene/alkane pathawy was only recently discovered and there has been no work so far to engineer the specific pathways that synthesize such hydrocarbons. Some species of algae are also known to naturally accumulate hydrocarbons that could serve as a fuel following chemical conversion. For example, certain races of the green alga *Botrococcus braunii* naturally secrete long-chain terpene hydrocarbons as a significant portion of their biomass (Eroglu and Melis, 2010; Metzger and Largeau, 2005). However, their use as a fuel source is made impossible by the incredibly slow growth rates of this alga (Cook et al., 2017). Other oleaginous algal species can accumulate a significant portion of their biomass as triacylglycerol compounds, generally under nitrogen stress. Indeed, this phenomenon drove the push for the use of algae as third generation biofuel feedstock in the first place. However, process design and downstream processing cost considerations of large-scale algal cultivation have hindered the common adoption of algal oils for transportation fuels (Quinn and Davis, 2015). Triacylglycerol stored by eukaryotic algae can also be turned into transportation fuels via transesterification to liberate the alkanes and alkenes from the glycerol backbone. An attractive alternative to the above concepts is instead to directly secrete ready-to-use hydrocarbon products from algal cells as this would overcome issues with biomass harvesting and chemical processing and thereby greatly reduce process costs (Delrue et al., 2013).

In order to achieve such a one-step conversion of CO_2_ into ready infrastructure-compatible hydrocarbons with photosynthetic hosts, however, genetic reprogramming becomes essential for introduction of synthetic metabolic pathways and optimization of the entire system. Several enzymes have recently been reported to enable biosynthesis of fatty aldehyde precursors (Akhtar et al., 2013), fatty alkanes (Bernard et al., 2012; Qiu et al., 2012), and fatty alkenes (Rude et al., 2011; Rui et al., 2015; Rui et al., 2014). Combinatorial assembly of such key enzymes into synthetic metabolic pathways consequently enabled a number of novel opportunities for hydrocarbon biosynthesis, as described by many including (Akhtar et al., 2013; Kallio et al., 2014; Sheppard et al., 2016; Zhu et al., 2017). Although such studies have so far only been reported using heterotrophic microorganisms *(Escherichia co¡i* and *Saccharomyces cerevisiae)* there are no reports of similar work in any phototrophic microorganism.

In this study, we describe a first and systematic study to implement synthetic metabolic pathways for the biosynthesis of hydrocarbon fuel in both prokaryotic and eukaryotic photosynthetic microorganisms using the model strains *Synechocystis* 6803 and *Chlamydomonas reinhardtii*. Several synthetic pathways towards saturated and unsaturated hydrocarbons were functionally demonstrated in *Synechocystis* 6803, increasing the hydrocarbon content up to 19-fold, and engineered *Chlamydomonas* accumulated 8-fold more alkenes than the wild-type. Interestingly, the “best” system was achieved by transferring a reconstructed pathway from eukaryotic algae into the prokaryotic cyanobacterium.

## MATERIALS AND METHODS

### 2.1. *Growth conditions, genetic constructs, transformation and screening of Escherichia coli and Synechocystis sp*. PCC 6803

*Escherichia coli* DH5α was used to propagate all the plasmids used in this study. Strains were cultivated in lysogeny broth (LB) medium (LB Broth, Sigma Aldrich), 37 °C, 180 rpm, and supplemented with appropriate antibiotics (final concentration: carbenicillin 100 μg/ml, chloramphenicol 37 μg/ml, kanamycin 50 μg/ml, gentamicin 10 μg/ml, and erythromycin 200 μg/ml).

*Synechocystis* sp. PCC 6803, obtained from Prof. Klaas Hellingwerf (University of Amsterdam, Netherlands), was cultivated in BG11 medium without cobalt ((hereafter BG11-Co), as the metal was used as an inducer in most cultures. All media contained appropriate antibiotic(s) (final concentration: kanamycin 50 μg/ml, gentamicin 50 μg/ml, and erythromycin 20 μg/ml). Gentamicin was only used for selection on agar plates. Precultures inoculated from colonies on agar plates were grown in 6-well plates (5 ml). When the OD730 reached 3-4, the culture was transferred to a 100-ml Erlenmeyer flask and the OD730 was adjusted to 0.2 by adding BG11-Co medium to a final volume of 25 ml containing appropriate antibiotic(s). The cultivation was carried out for 10 days at 30 °C with continuous illumination at 60 μmol photons m^−2^ s^−1^ and 1% (v/v) CO_2_. Each main treatment culture was induced on day 2 and samples were taken for measurement of OD730 and metabolites day 6 and 10. All cultivations were carried out in an AlgaeTron230 (Photon Systems Instruments) (PSI) at 30 °C with continuous illumination at 60 μmol photons m^−2^ s^−1^ and 1% (v/v) CO_2_, except where noted (100-300 μmol photons m^−2^ s^−1^). A representative growth curve and all final OD730 values are shown in Supplementary Figure 1.

All plasmids (Supplementary Table 1A) used for transformation of cyanobacteria were assembled using the BASIC Assembly method (Storch et al., 2015). Linkers were designed using the R20DNA software: http://www.r2odna.com/ and obtained from Integrated DNA Technologies Incorporated. The details of all linkers, primers and DNA parts used to construct each plasmid are given in Supplementary Tables 1B, 1C and 1D.

For transformation by natural assimilation, each *Synechocystis* sp. PCC 6803 strain was inoculated from freshly prepared colonies on agar plates into 25 ml BG11-Co with a starting OD 0.02. The cells were harvested when the OD730 reached 0.4-0.7, washed in 10 ml BG11-Co twice, and resuspended in 500 μL BG11-Co. One hundred microliters of concentrated liquid culture were mixed with four to seven micrograms of plasmid and incubated at 30°C with continuous illumination at 60 μmol photons m^−2^ s^−1^ and 1% (v/v) CO_2_ for 12-16 h prior to plating on BG11-Co agar containing 10% strength of antibiotic. To promote segregation, individual colonies were restreaked on BG11-Co agar with higher antibiotic concentration. To check the segregation, the biomass was resuspended in nuclease free water and exposed to two freeze-thaw cycles (95°C, −80°C). Following centrifugation, 3 μL was used as a template for a diagnostic polymerase chain reaction (PCR). Primers used for each PCR are listed in Supplementary Table 1C. Only fully segregated mutants were used in further experiments. All cyanobacteria strains used in the study are listed in Supplementary Table 2.

For transformation by triparental conjugation, one hundred microliters of the cargo strain (*E. coli* HB101 (already carrying the pRL623 plasmid)), conjugate strain (ED8654 (Elhai and Wolk, 1988)), and *Synechocystis* sp. PCC 6803 (OD730 ~1) were mixed and incubated for 2 h (30 °C, 60 μmol photons m^−2^ s^−1^). Prior to mixing, all the *E. coli* and cyanobacteria strains were washed with fresh LB and BG11-Co medium, respectively, to remove the antibiotics. After 2 h of incubation, the culture mix was transferred onto BG11 agar plates without antibiotic and incubated for 2 d (30 °C, 60 μmol photons m^−2^ s^−1^). After 2 d of incubation, cells were scraped from the agar plate, resuspended in 500 μL of BG11-Co medium, and transferred onto a new agar plate containing 20 μg/ml erythromycin. Cells were allowed to grow for one week until colonies appeared. Individual colonies were restreaked onto a new plate containing 20 μg/ml erythromycin and used for subsequent experiments.

### 2.2. Growth conditions, genetic constructs, transformation and screening of Chlamydomonas reinhardtii

*C. reinhardtii* strain UVM4 was used in this work (Neupert et al., 2009) graciously provided by Prof. Dr. Ralph Bock)). The strain was routinely maintained on Tris acetate phosphate (TAP) medium (Gorman and Levine, 1965) either with 1.5% agar plates or in liquid with 250 μmol photons m^−2^ s^−2^. Transformation was conducted with glass bead agitation as previously described (Kindle, 1990). The amino acid sequences of *C. reinhardtii* native fatty acid photodecarboxylase (FAP) (Uniprot: A8JHB7; (Sorigué et al., 2017)), *E. coli* thioesterase A (TesA: P0ADA1), *Jeotgalicoccus* sp. ATCC 8456 terminal olefin-forming fatty acid decarboxylase (OleTJE) (E9NSU2), and *Rhodococcus sp*. NCIMB 9784 P450 reductase RhFRED (Q8KU27) were codon optimized and copies of the intron 1 of ribulose bisphosphate carboxylase small subunit 2 (RBCS2) were added throughout the coding sequences as previously described (Baier et al., 2018). The nucleotide sequences of optimized intron containing genes have been submitted to NCBI, accession numbers can be found in Supplementary Table 3. All synthetic genes were chemically synthesized (GeneArt) and cloned between SamHI-SglII in the pOpt2_PsaD_mVenus_Paro or pOpt2_PsaD_mRuby2_Ble vectors (Wichmann et al., 2018). PsaD represents the 36 amino acid photosystem I reaction center subunit II (PsaD) chloroplast targeting peptide (CTP) (Lauersen et al., 2015) between NdeI-BamHI restriction sites of the pOpt2 vectors (Wichmann et al., 2018). The native FAP enzyme was designed to contain an additional glycine codon at aa position 33 to allow the insertion of a BamHI site at the border of the predicted CTP. The whole synthetic enzyme including native targeting peptide was cloned *NdeI-BglII* and a version was created with the PsaD CTP built by cloning BamHI-SglII into the vectors described above. Fusions of different sequences were made by digestion and complementary overhang annealing of the BamHI-*Sg*lII mediated restriction sites for each respective construct as needed to obtain the fusions used in the present work (Supplementary Figure 2). After transformation, expression was confirmed by fluorescence microscopy screening for mVenus (YFP) or mRuby2 (RFP) reporters as previously described (Lauersen et al., 2016; Wichmann et al., 2018). Individual mutants were subjected to Western blotting and immuno detection to determine whether full-length protein products were formed (anti-GFP polyclonal HRP linked antibody, Thermo Fisher Scientific). Wide-field fluorescence microscopy was used to confirm chloroplast localization of YFP-linked constructs as previously described (Lauersen et al., 2016).

### 2.3. Product analysis

Three different extraction and analysis protocols were used for the analysis of (1) acids, (2) alcohols and (3) alkanes as well as alkenes from cyanobacteria cultures. For each analyte group, liquid cultures in flasks were mixed well by shaking prior to transferring 2 mL of liquid culture into a PYREX round bottom threaded culture tube (Corning, Manufacturer Part Number: 99449-13).

For fatty acid analysis, free fatty acid extraction was performed as described previously (Liu et al., 2011; Yunus and Jones, 2018). In brief, two hundred microliters of 1 M H3PO4 were added to acidify each 2 mL culture and spiked with 100 μg pentadecanoic acid (Sigma Aldrich) as an internal standard. Four millilitres of n-hexane (VWR Chemicals) was added and the mixture vortexed vigorously prior to centrifugation at 3500 x g for 3 min. The upper hexane layer was then transferred to a fresh PYREX round bottom threaded culture tube and evaporated completely under a stream of nitrogen gas. Five hundred microliters of 1.25 M HCl in methanolic solution were added to methyl esterify the free fatty acid at 85 °C for 2 h. Samples were cooled to room temperature and 500 μL of hexane was added for extraction of the fatty acid methyl esters (FAMEs).

For fatty alcohol, alkane and alkene analysis, extraction was done as described previously (Zhou et al., 2016) with modification. Briefly, 2 mL of liquid culture were spiked with 50 μg 1-nonanol, 100 μg octadecane, and 100 μg 1-pentadecanol and mixed with 4 mL of chloroform:methanol (2:1 v/v) solution. The mixture was vortexed vigorously and centrifuged at 3500 x g for 3 min. The lower organic phase was then transferred into a new glass tube and extraction was repeated one more time. The lower organic phase was combined and dried under a stream of nitrogen gas. For fatty alcohol derivatisation, the dried extract was resuspended in 100 μL chloroform, mixed with 100 μL of N, O-bistrifluoroacetamide (BSTFA) (TCI Chemicals) and transferred to an insert in a GC vial that was incubated at 60 °C for 1 h prior to GC analysis. Note that no derivatisation was needed for the analysis of hydrocarbons.

Samples (1 μL) were analysed using an Agilent Technologies (Santa Clara, CA, USA) 7890B Series Gas Chromatograph (GC) equipped with an HP-5MS column (pulsed split ratio 10:1 and split flow 10 ml/min), a 5988B Mass Spectrophotometer (MS) and a 7693 Autosampler. For the acids the GC oven program followed an initial hold at 40 °C for 3 min, a ramp at 10 °C.min^−1^ to 150 °C, a second ramp at 3 °C.min^−1^ to 270 °C, a third ramp at 30 °C.min^−1^ to 300 °C, and a final hold for 5 min. For alcohols and alkenes, there was an initial hold at 40 °C for 0.5 min, a ramp at 10 °C.min^−1^ to 300 °C, and a final hold for 4 min. For alkanes, the oven was initially held at 70 °C for 0.5 min, a ramp at 30 °C.min^−1^ to 250 °C, a second ramp at 40 °C.min^−1^ to 300 °C, and a final hold for 2 min. The acids, alkanes and alcohols were quantified by comparing the peak areas with that of the internal standards: pentadecanoate (for all acids), octadecane (for all alkanes), 1-nonanol (for C8 to C12 alcohols) and 1-pentadecanol (for C14 alcohols and above). The quantity of the main products (C15 and C17 alkanes, C15 alkene, and C12, C14, C16, and C18 alcohols and acids) were also corrected with their respective mass spectrometer response factors obtained using dilution series of commercial standards.

Gas chromatography mass spectroscopy (GC-MS) aimed at identification of hydrocarbon products from *C. reinhardtii* was conducted with solvent extracted samples following previously described protocols and internal standards (Lauersen et al., 2016). Quantification of 7-heptadecene was performed with serial dilutions (1 to 900 μM) of commercial 1-heptadecene standard (Acros Organics) in dodecane using extracted ion chromatograms with masses 55.00, 69.00, 91.00, 93.00, 83.00, 97.00, and 111.00.

## RESULTS AND DISCUSSION

Several synthetic pathway designs were considered, all commencing with the liberation of “free” fatty acids from the native fatty acid biosynthesis pathway (Fig. 1), the presumed native precursor for many of the decarboxylating enzymes evaluated in this study.

**Figure 1.**
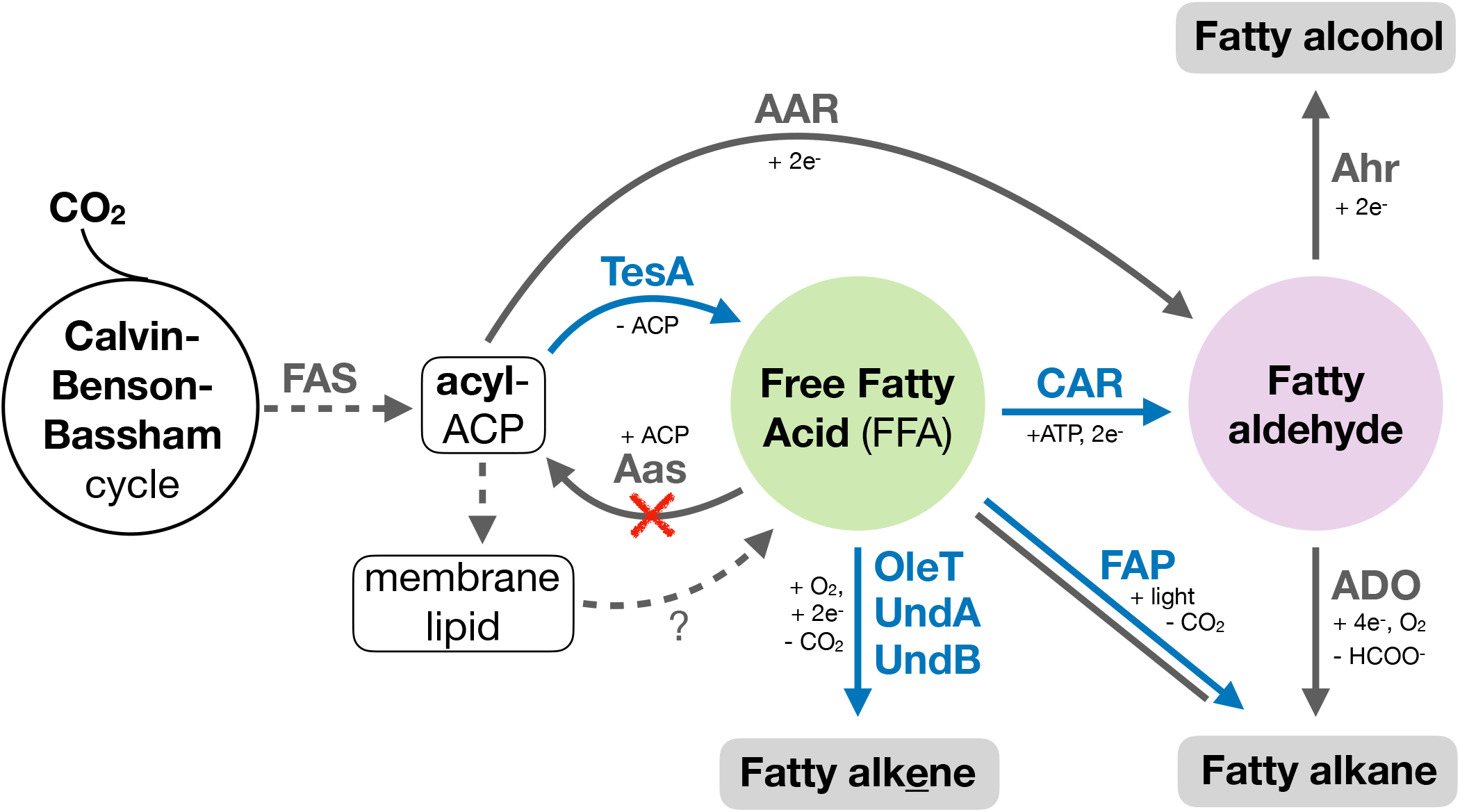
Native and synthetic metabolic pathways evaluated in the present study with incomplete stoichiometry. The graphic illustration shows the introduced TesA (thioesterase (Cho and Cronan, 1995)), CAR (carboxylic acid reductase (Akhtar et al., 2013)), UndA (responsible for 1-undecene biosynthesis in *Pseudomonas* (Rui et al., 2014)), UndB (also responsible for 1-undecene biosynthesis in *Pseudomonas* (Rui et al., 2015)), OleT (responsible for olefin biosynthesis in *Jeotgalicoccus* (Rude et al., 2011)) and FAP (fatty acid photodecarboxylase (Sorigué et al., 2017)) enzymes alongside the native AAR/ADO (acyl-ACP reductase and aldehyde deformylating oxygenase (Schirmer et al., 2010)), AHR (aldehyde reductase, unknown) and FAP enzymes. Blue reactions are non-native and those in grey are native. The red cross indicates deletion of the *aas* gene.

### 3.1. Over-production of free fatty acids as precursor for hydrocarbon biosynthesis - Expression of Escherichia thioesterase deregulates lipid membrane biosynthesis in Chlamydomonas

In order to liberate FFAs in cyanobacteria we over-expressed the *E. coli* C16-C18 specific thioesterase TesA (Cho and Cronan, 1995) lacking its native signal sequence peptide (‘TesA) and deleted the gene encoding the native fatty acyl ACP synthase (aas) (Kaczmarzyk et al., 2010; Liu et al., 2011)(Fig. 1). The native signal sequence peptide directs TesA to the periplasm in *E. coli* (Cho et al 1993) and its removal is assumed to maximize the liberation of “free” fatty acids also in cyanobacteria by retaining the enzyme in the cytosol. Such ‘TesA/Δaas engineering has previously been reported several times before in cyanobacteria (Liu et al., 2011, Ruffing et al., 2014; Work et al., 2015; Kato et al., 2017), with 13% (g/g cell dry weight (CDW)) as the highest reported fatty acid yield in *Synechocystis* 6803 (Liu et al 2011). Further potentially stackable modifications to the strain or process have also been reported. For example, by employing a solvent overlay, Kato et al., 2017 reported up to 36% (g/g) cdw of fatty acids excreted into the media using ‘TesA/Δ*aas Synechococcus elongatus sp*. PCC 7942. In the present study, the chromosomal integration of ‘*tesA* into the *psbA2* site (slr1311) of *Synechocystis* 6803 Δ*aas* (Δ*aas*-’TesA), under the control of the light-inducible promoter PpsbA2S, resulted in the excretion of of C14:0 (3.5 mg/g CDW), C16:0 (23.2 mg/g CDW) and C18:0 (5.7 mg/g CDW) fatty acids with a chain-length distribution that is in agreement with previously reported findings (Liu2011) (Fig. 2A; Supplementary Fig. 3).

**Figure 2.**
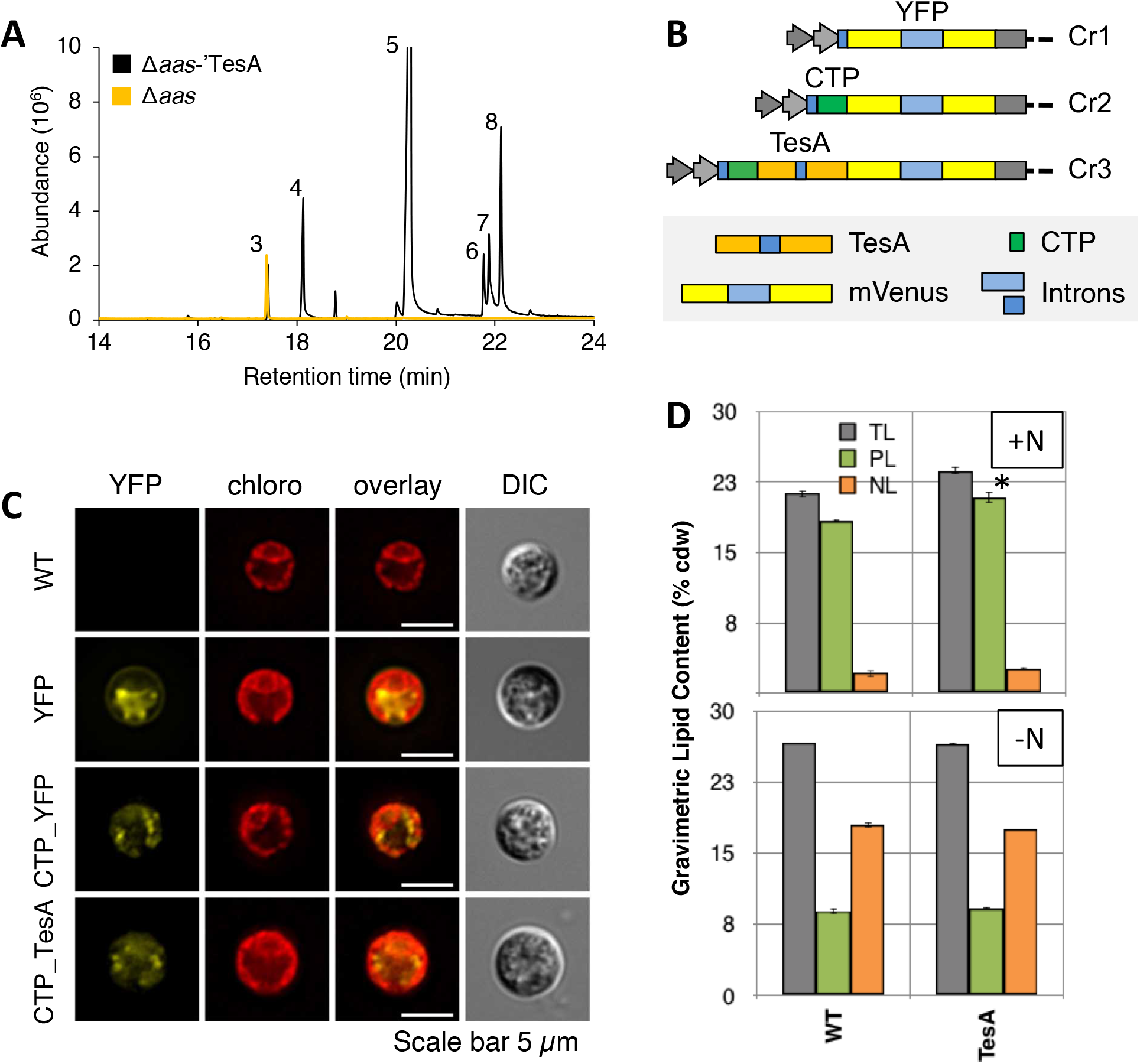
Engineering for enhanced accumulation of free fatty acids. (A) Representative total ion count chromatograms for *Synechocystis* 6803 strains Δ*aas*-’TesA (black) vs. Δ*aas* only (orange) extracted on day 10 of cultivation (induced day 2). Peak identities: (3) Heptadecane, (4) Tetradecanoic acid, (5) Hexadecanoic acid, (6) 9,12-octadecadienoic acid, (7) 9-octadecenoic acid, (8) Octadecanoic acid. (B) Graphic representation of the constructs used to transform *Chlamydomonas*. CTP = Chloroplast Transit Peptide. (C) Fluorescence microscopy of representative strains indicating appropriate chloroplast localization of the CTP_’TesA_YFP construct. (D) Total (TL), polar (PL), and neutral (NL) gravimetric lipid fractions of *Chlamydomonas* parental strain and TesA overproducing strains under nutrient replete conditions (N+) and after 96 hours of nitrogen depletion (N-). PL is significantly greater in +N for TesA: ttest, p:0.047 (indicated by an asterisk).

Overproduction of the same thioesterase (‘TesA) and targeting of the enzyme product to the chloroplast was possible in *C. reinhardtii*. The synthetic algal optimized *E. coli* ‘*tesA* gene was fused with an N-terminal PsaD-based chloroplast targeting peptide and a C-terminal yellow fluorescent protein (YFP) encoding gene. Both the coding genes were interspersed by synthetic introns (Fig. 2B) as previously described to enhance transgene expression from the nuclear genome (Baier et al., 2018). Fluorescence microscopy indicated correct localization of the ‘TesA fluorescent protein fusion to the algal chloroplast (Fig. 2C). Although no FFA could be detected in the culture medium, a difference was observed in the lipid profile of the green algal cells, suggesting a de-regulation of fatty acid synthesis that specifically affected the polar lipid fraction of the alga. This was indicated by an over-accumulation of C18:1n9c chain lengths in the polar lipid membranes, with subtle changes observed in other acyl-ACP species such as C14:0 (Fig. 2D; Supplementary Fig. 4). Thus, ‘TesA_YFP clearly had an impact on lipid metabolism in the eukaryotic algal host, but, the capture of liberated FFA by acyl-ACP or -CoA synthases is likely too effective, thereby limiting the application of the same engineering principles carried out for cyanobacteria. An annotated gene product in *Chlamydomonas* Cre06.g299800 (Phytozome v5.5) has some sequence similarity to *Synechocystis aas* and therefore represents an interesting target for future strategies to block native re-uptake of FFA in the green algal cell.

Having achieved strains with enhanced accumulation of FFA in *Synechocystis*, or at least a perturbation to the lipid biosynthetic system in *Chlamydomonas*, we proceeded to investigate enzymes that further convert FFAs into hydrocarbon end-products.

### 3.2. Effective conversion of free fatty acids into alkenes using UndB

Three different enzymes that catalyze the conversion of fatty acids into alkenes have been recently reported, OleT (Rude et al., 2011), UndA (Rui et al., 2014), and UndB (Rui et al., 2015) (Fig. 1). So far, the best reported productivity in both *E. coli* (Rui et al., 2015) and *S. cerevisiae* (Zhou et al., 2018) has been with UndB.

In *Synechocystis* 6803, we transformed the Δ*aas*-’TesA strain with an RSF1010-based plasmid harboring a codon-optimized *undB* under the control of the Pclac143 promoter (Markeley et al., 2014), thereby generating the strain Δ*aas*-’TesA-1010-UndB (Fig. 3A). After 10 days of cultivation, both the free fatty acids and alkanes were extracted and analyzed as described in the Materials and Methods section. The accumulation of free fatty acids was markedly reduced in the Δ*aas*-’TesA-1010-UndB strain (Fig. 3B, 3C). In its place, both 1-pentadecene and 1-heptadecene accumulated with a molar yield suggesting approximately 55% conversion of ‘TesA-liberated FFAs (compare Fig. 3C with Fig. 3D). More than >84% of the FFAs disappeared relative to the Δ*aas*-’TesA strain suggesting that UndB was catalytically efficient *in vivo* and that the electrons required in the UndB reaction were fortunately supplied by an unknown source. The Δ*aas*-’TesA-1010-UndB strain displayed a lower biomass accumulation than the controls (Δ*aas*-empty and Δ*aas*-’TesA strains) (Supplementary Fig. 1), presumably due to product toxicity imparted by the alkenes. A direct comparison with the conversion efficiency in *E. coli* is not possible since the FFA conversion efficiency was not reported in the original work (Rui et al., 2015). Despite the disappearance of C14:0 fatty acids in the Δ*aas*-’TesA-1010-UndB strain, no measurable 1-tridecene (the expected corresponding alkene) was observed in the whole culture extracts (Fig. 3C). None of the observed alkene products were secreted extracellularly (Fig. 3E).

**Figure 3.**
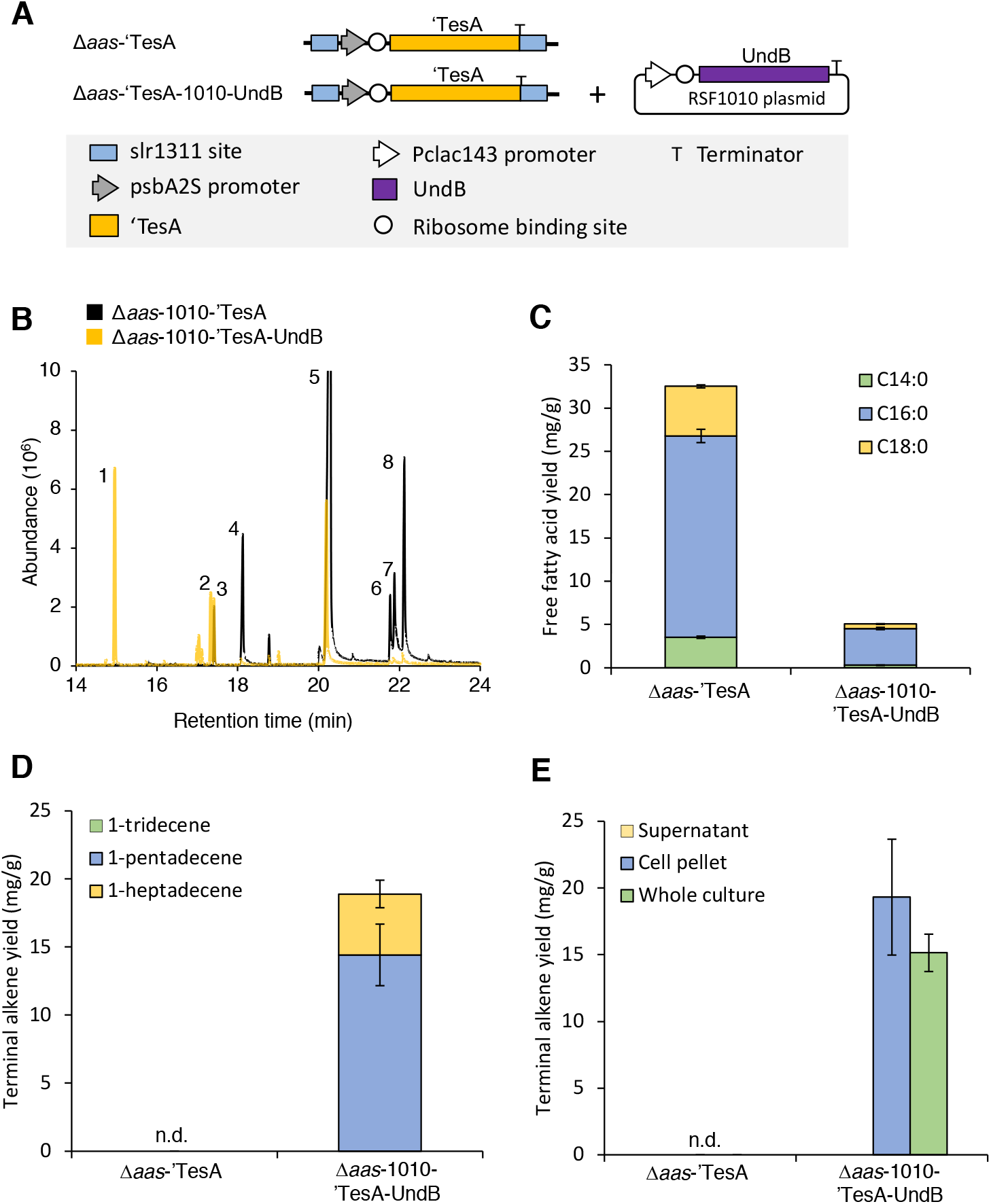
Over-expression of UndB results in effective (>50%) conversion of fatty acids into corresponding alkenes. (A) Graphic representation of the genetic modification of *Synechocystis* sp. PCC 6803 and the plasmid used for UndB expression. (B) GC-MS chromatograms with extracts from the two different strains (w/wo UndB); Δ*aas*-1010-’TesA (black) and Δ*aas*-1010-’TesA-UndB (orange). (C) The free fatty acid yield (relative to biomass) in the whole cultures of the two strains, subdivided into the three dominant chain-lengths. (D) The yield of alkenes in the whole cultures of the two strains, subdivided into the three dominant chain-lengths. (E) The localization of the alkene products in whole cultures of the two strains. Peak identities: (1) 1-pentadecene, (2) 1-heptadecene, (3) heptadecane, (4) tetradecanoic acid, (5) hexadecanoic acid, (6) 9,12-octadecadienoic acid, (7) 9-octadecenoic acid, (8) octadecanoic acid. Data are mean ± SD from three biological replicates. All samples were extracted on day 10.

In *Chlamydomonas*, we attempted to over-produce the *Jeotgalicoccus* sp. terminal olefin-forming fatty acid decarboxylase (OleTJE) and the *Rhodococcus sp*. P450 reductase (RhFRED). OleTJE was chosen as it could theoretically produce C17:1 and C15:0 hydrocarbons from the major lipid species of the green algal cell, C18:1 and C16:0, respectively (Fig. 1). Fusion to RhFRED has been reported to enable hydrogen peroxide-independent decarboxylase activity (Liu et al., 2014). The protein products of this decarboxylase and its fusion in either orientation to RhFRED could be detected by Western blotting and located to the algal chloroplast in fluorescence microscopy (Supplementary Figure 5). However, no differences in GC-MS profiles between the parental and expression strains could be found in either dodecane solvent overlays or cell-pellet solvent extracts.

### 3.3. Transfer of the CAR/ADO based pathway from E. coli to Synechocystis 6803 resulted in the accumulation of fatty alcohols and a reduction in alkane accumulation

Carboxylic acid reductases (CAR) have been previously used to construct a number of synthetic pathways for alkane biosynthesis in heterotrophic microorganisms (Akhtar et al., 2013; Kallio et al., 2014; Sheppard et al., 2016; Zhu et al., 2016). Although CAR appears to have a high capability for converting fatty acids into corresponding fatty aldehydes (Akhtar et al., 2013) (Fig. 1), a bottleneck in previous heterotrophic pathways is the subsequent conversion into alkanes by kinetically slow ADO enzymes and competition with native aldehyde reductases that more effectively convert aldehydes into alcohols (Kallio et al., 2014; Sheppard et al., 2016).

Since *Synechocystis* 6803 natively harbors an aldehyde deformylating oxygenase (ADO) with the appropriate substrate specificity (Khara et al., 2013) (Fig. 1), we first combined TesA with CAR and evaluated its ability to supply the native ADO. A synthetic operon expressing all required parts (including the CAR maturation protein Sfp) was introduced to the RSF1010 plasmid backbone (Fig. 4A) and used to transform *Synechocystis* 6803 *Δaas*, thus creating the strain Δ*aas*-1010-TPC2. This strain accumulated both fatty acids (Fig. 4B and 4D) and fatty alcohols (Fig. 4C and 4E). The quantity of heptadecane was reduced in Δ*aas*-1010-TPC2 relative to Δ*aas*-1010-’TesA (Fig. 4F). This suggests that the introduced CAR-based pathway had not managed to increase the supply of fatty aldehydes to the native ADO. CAR and native aldehyde reductase(s) had instead very effectively converted >90% of the FFA pool (Fig. 4D) into corresponding alcohols (Fig. 4E). The most likely reason for the increase in FFA in latter experiments is due to increased expression of ‘TesA using the RSF1010 plasmid in Δ*aas*-1010-‘TesA (Fig. 4D), relative to the amount of ‘TesA when expressed from the chromosomal location in Δ*aas*-’TesA (Fig. 3C). Similar observations have also been previously reported by Angermayr et al. (Angermayr et al., 2014). The different promoters used in the two strains are also likely to have influenced the outcome, however, we are not aware of any studies that directly compare the two promoters head-to-head.

**Figure 4.**
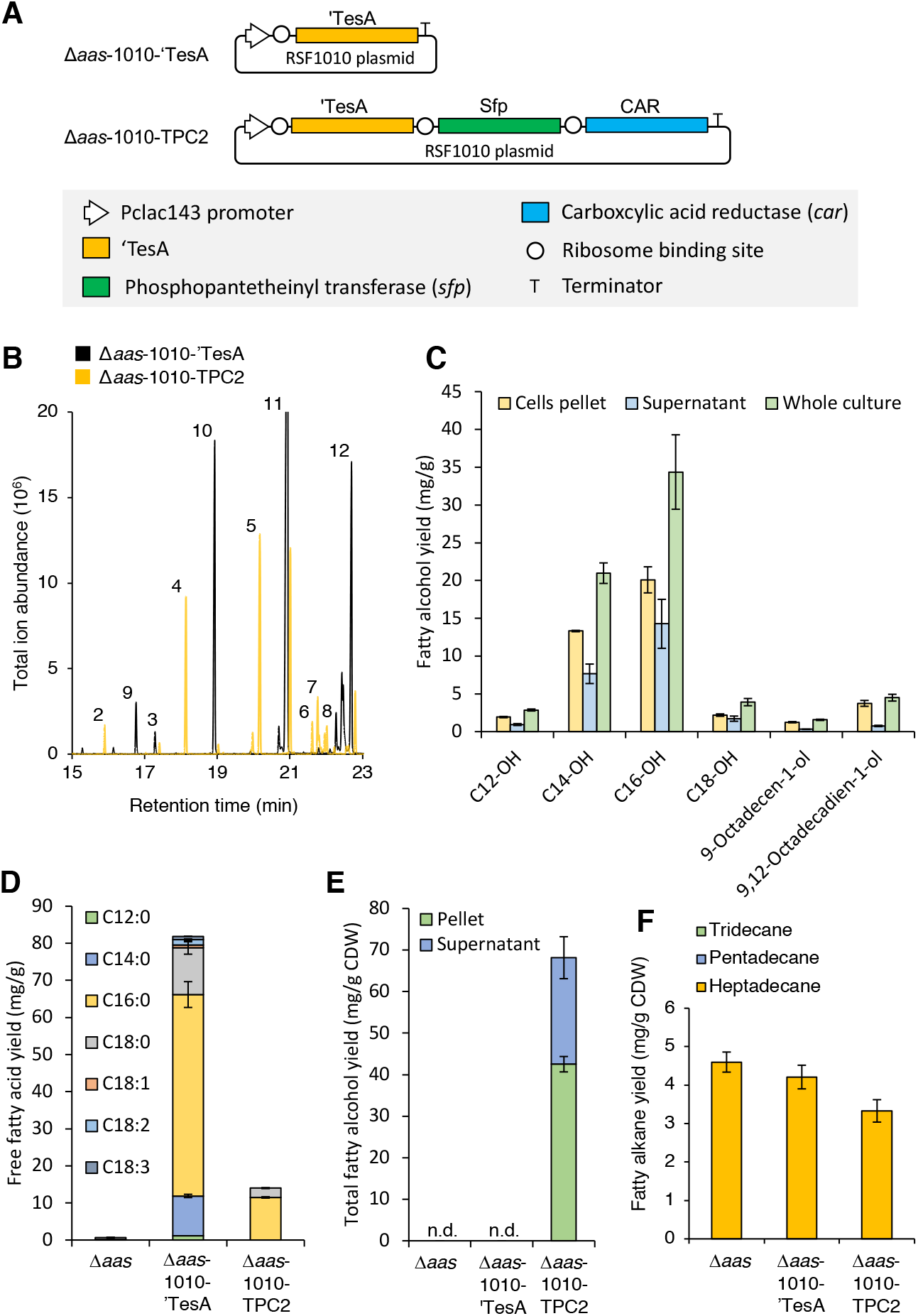
The CAR-dependent pathway produces mainly fatty alcohols. (A) Graphic overview (not to scale) illustrating the main constructs studied in the figure. (B) Total ion chromatogram from extracts of Δ*aas*-1010-’TesA (black) and Δ*aas*-1010-TPC2 (orange). (C) Fatty alcohol profile from extracts of Δ*aas*-TPC2. The yield of fatty acids (D), alcohols (E) and alkanes (F). Peak identities: (2) 1-dodecanol, (3) heptadecane, (4) 1-tetradecanol, (5) 1-hexadecanol, (6) 9,12-octadecadien-1-ol, (7) 9-octadecen-1-ol, (8) 1-octadecanol, (9) dodecanoic acid, (10) tetradecanoic acid, (11) hexadecanoic acid, (12) octadecanoic acid. Data are mean ± SD of three biological replicates. Cultures were induced on day 2 following dilution and samples were extracted on day 10.

Substantial quantities of fatty alcohols did accumulate in the Δ*aas*-1010-TPC2 strain, suggesting that the supply of fatty aldehydes is not the limiting factor. One possibility is that the native aldehyde reductases are simply much more active than the native ADO (Eser et al., 2011; Lin et al., 2013). Another possibility is that native ADO and AAR form a close metabolon *in vivo* (Warui et al., 2015) that locks out access to ADO from external supplies of fatty aldehydes. In order to test this possibility, we attempted to create a variant of Δ*aas*-1010-TPC2 that also included chromosomal ADO over-expression casette under the PpsbA2S promoter. Despite numerous transformation and segregation attempts, however, we were unable to isolate any stable segregants. Another complementary strategy that could be considered in future work would be to eliminate native aldehyde reductases, as previously carried out in earlier *E. coli* studies (Kallio et al., 2014; Sheppard et al., 2016), although the full complement of fatty aldehyde reductase encoding genes in cyanobacteria remains unknown. Given the lack of success in producing alkanes with the CAR/ADO route in cyanobacteria we then considered alternative options for both cyanobacteria and algae.

### 3.4 Engineering of the native eukaryotic algae pathway and transfer to cyanobacteria results in enhanced conversion of CO_2_ into alkanes

A fatty acid photodecarboxylase (FAP) that directly converts saturated and unsaturated FFAs into alkanes and alkenes, respectively, was recently discovered in eukaryotic algae (Sorigué et al., 2017). In *Chlamydomonas*, the source of free fatty acids for the native alkene pathway remains unknown, although the degradation of membrane lipids may release some FFA (illustrated in Fig. 1). However, we would expect increased accumulation of alkanes in algae if we were able to increase the cellular quantity of the native FAP and/or introduce synthetic routes to the FFA precursors.

Accordingly, we overproduced native FAP from *C. reinhardtii* (CrFAP) on its own or in combination with co-production of *E. coli* ‘TesA. The over-expression of CrFAP was carried out either with its native chloroplast targeting peptide (CTP) or the robust PsaD CTP which has been previously used to mediate chloroplast localization of numerous reporters (Lauersen et al., 2015; Lauersen et al., 2018; Rasala et al., 2013). In order to minimize any native regulation of the genomic sequence, the gene was subjected to a strategy of gene design which has recently been shown to enable robust transgene expression from the nuclear genome of this alga (Baier et al., 2018). Briefly, the sequence was codon optimized based on its amino acid sequence and multiple copies of the first intron of the *C. reinhardtii* ribulose-1,5-bisphosphate carboxylase/oxygenase (RuBisCo) small subunit 2 (rbcS2i1, NCBI: X04472.1) were spread throughout the coding sequence *in silico*. This nucleotide sequence was chemically synthesized and used for expression from the algal nuclear genome. This strategy has previously enabled heterologous overproduction of non-native sesquiterpene synthases (Lauersen et al., 2016; Lauersen et al., 2018; Wichmann et al., 2018), and in the present study also the ‘TesA, OleTJE, and RhFRED proteins. However, complete codon optimization and synthetic intron spreading of a native gene has not yet been demonstrated in eukaryotic algae. Both constructs mediated full-length target protein production which was detectible in Western blots (Supplementary Fig. 5B). Replacing the native CTP with the PsaD CTP enabled more reliable and robust accumulation, which was detectible as YFP signal in the algal chloroplast (Supplementary Fig. 6) and strong bands in transformants expressing this construct in Western blots (Supplementary Fig. 5B). The parental UVM4 strain was found to contain ~0.5 mg/g 7-heptadecene as a natural product (Supplementary Fig. 7). Transformants generated with the CrFAP construct (Cr8) were found to contain up to 8x more of this alkene compared to the empty vector (Cr2) control strain (up to 8.5 ± 1.5 mg/g, Fig. 5) which was found almost exclusively within the biomass (Supplementary Fig. 7). The product was not detected in dodecane solvent overlays. CrFAP accepts a very specific substrate (cis-vaccenic acid, C18:1cisΔ11) *in vivo* (Sorigué et al., 2017), which corresponds to the accumulation of only 7-heptadecene as the only detected increased product. This substrate is an unusual FA, and is likely not naturally abundant in the algal cell. Notably, any attempts to increase the avaialability of free fatty acids using *E. coli* ‘TesA did not result in any increase in the quantity or diversity of accumulated alkanes. Future enzyme engineering will likely be able to overcome this substrate specificity and increase overall yields of liberated hydrocarbons. However, a strategy which would allow secretion of these molecules, similar to the capture of heterologous terpenoids in dodecane solvent overlay (Lauersen et al., 2016; Lauersen et al., 2018; Wichmann et al., 2018), would be an attractive next target in order to enable photo-biocatalysis of hydrocarbons from the algal biomass.

**Figure 5.**
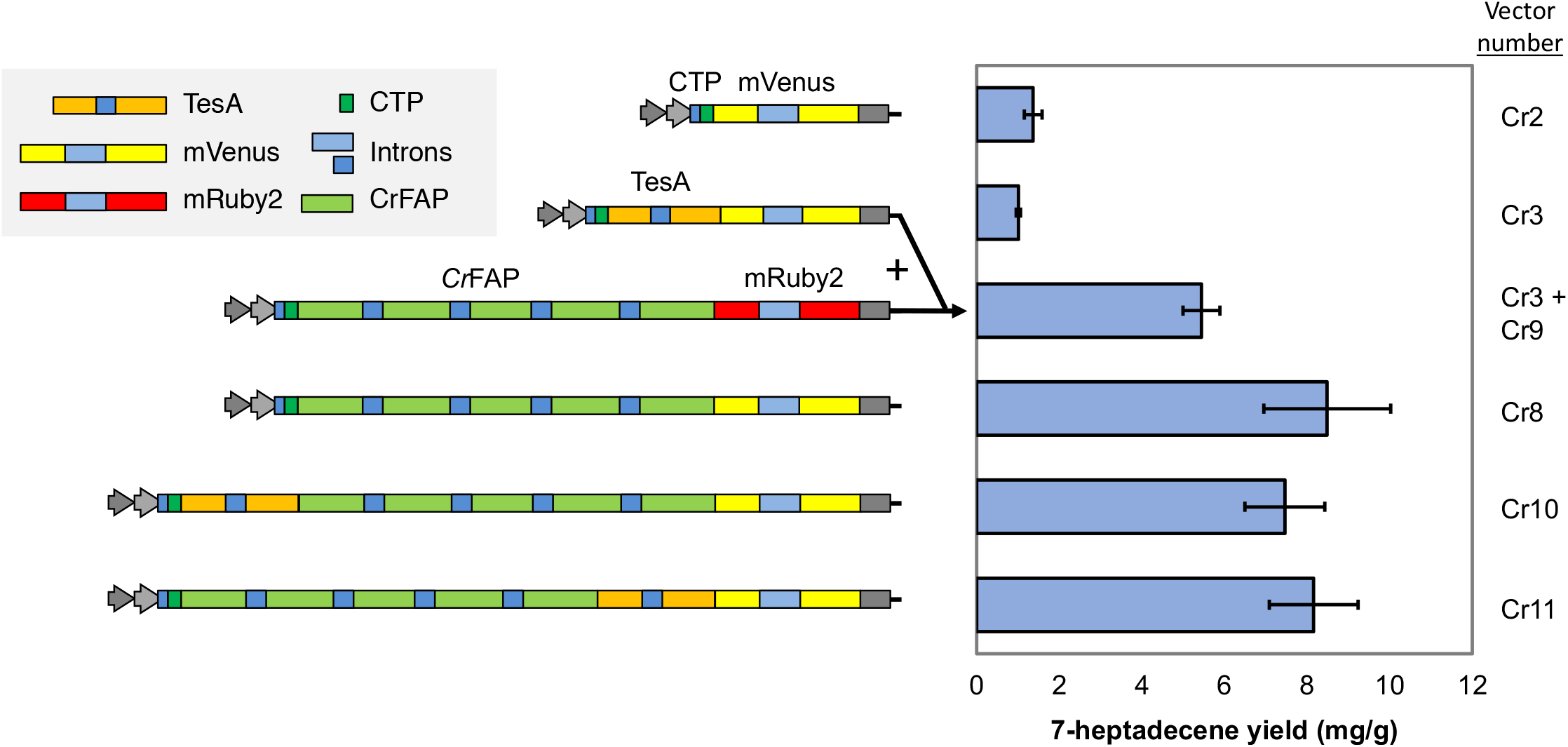
CrFAP over-expression increases 7-heptadecene yield, but heterologous thioesterase (TesA) expression, its co-expression, and C-or N-terminal fusion with CrFAP has no benefit. Mutants expressing indicated constructs (left panel) were cultivated for seven days in TAP medium with 250 μmol photons s^−1^ m^−2^ constant illumination and cell pellets were extracted with cell rupture by glass beads and dodecane for yield quantification of 7-heptadecene via GC-MS (bar graph, right). All constructs bear a PsaD chloroplast targeting peptide (CTP) to allow protein transit to the chloroplast. Arrows and plus sign indicate co-expression in double transformed mutants. Error bars represent 95% confidence intervals of single strains cultivated in biological triplicates.

Given the success with the FAP pathway in *Chlamydomonas* (present study) and earlier work in *E. coli* (Sorigué et al., 2017), as well as finding that ‘TesA expression can substantially enhance the FFA pool in cyanobacteria, a synthetic FAP pathway was an obvious choice to consider also for the prokaryotic host. We therefore proceeded to implement a reconstituted variant of the eukaryotic algae pathway in cyanobacteria by combining TesA with FAP. Given the genetic instability challenges with the CAR/ADO system (see Section 3.3) we shifted our constructs to the more tightly repressed Pcoa promoter (Peca et al., 2008) for controlling the expression of *E. coli* TesA and the *Chlorella variabilis* FAP from the RSF1010 plasmid (Fig. 6A). We noted that the yield of FFA was substantially increased when driving the expression of TesA with the Pcoa promoter (Fig. 6C) compared to Pclac143 (Fig. 4D).

**Figure 6.**
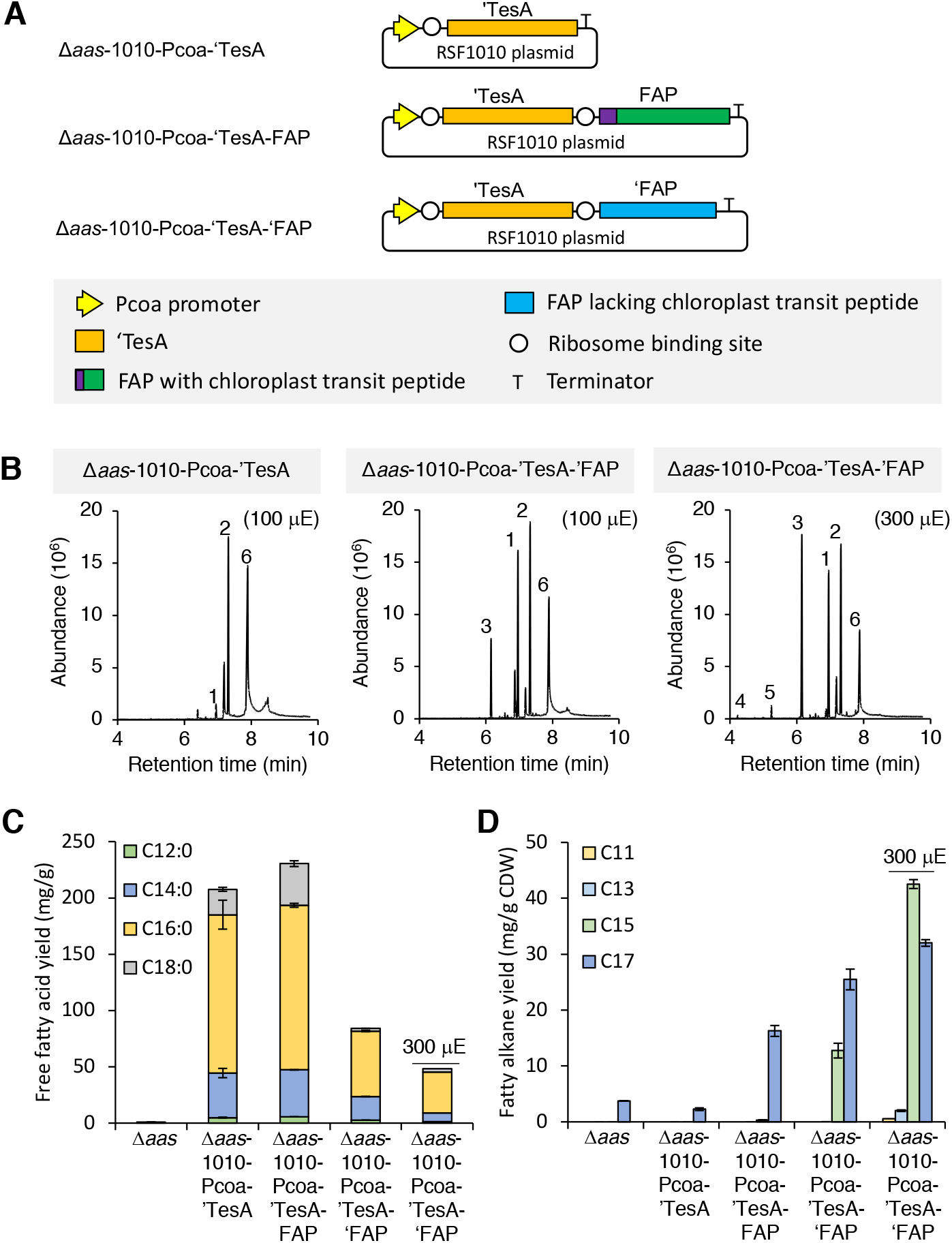
Conversion of free fatty acids into alkanes in cyanobacteria using FAP. (A) Graphic representative of the plasmids used to transform *Synechocystis* sp. PCC 6803. (B) Total ion chromatogram from Δ*aas*-Pcoa-’TesA (left) and Δ*aas*-1010-Pcoa-’TesA-’FAP (100 mE, middle; 300 mE, right). The free fatty acid (C) and alkane (D) yield in all tested strains. Data are mean ± SD of three biological replicates. Samples were extracted on day 10. Peak: (1) heptadecane, (2) octadecane (internal standard), (3) pentadecane, (4) undecane, (5) tridecane, (6) hexadecanoic acid.

Despite the dominance of C16:0 fatty acids released by ‘TesA in *Synechocystics* 6803, alongside minor fractions of C14:0 and C18:0, the C17:0 alkanes dominated the hydrocarbon fraction at the lower light intensity (100 μE) (Fig. 6B and 6D). This alkane profile in *Synechocystis* 6803 is very different to that observed in *E. coli* without over-expression of ‘TesA (see Fig. S4 in (Sorigué et al., 2017)). We also observed substantial peaks of 8-heptadecene and 6,9-heptadecadiene, as suggested by comparison with a NIST mass spectrometry library, although a lack of standards prohibited confirmation (Supplementary Figure 8). Curiously, these alkenes were only detected at day 6 and were not present in samples harvested on day 10. As the fatty chain-length profiles differ when the same thioesterase is expressed in different *E. coli* strains (Akhtar et al., 2015; Jing et al., 2011), this suggests that the *in vivo* product profile of any thioesterase-dependent pathway also is dependent on what the fatty acid synthesis pathway provides, not just the substrate specificity of the thioesterase used.

Removal of the predicted chloroplast targeting sequence of FAP (‘FAP) resulted in a doubling of the alkane yield, this time accompanied also by C15 pentadecane. As the FAP reaction is light-dependent, we also did a simple evaluation of this environmental factor. When the light intensity was tripled, the total alkane production with the Δ*aas*-1010-’TesA-‘FAP strain increased to a yield of 77.1 mg/g CDW (19-fold enhancement relative to Δ*aas*) and a titer of 111.2 mg/L. The product profile also shifted (Fig. 6D) despite the lack of a similar shift in the remaining FFA fraction (Fig. 6B), suggesting that the substrate specficity of FAP is flexible and interestingly might change in response to a change in its cellular environment.

At 100 μE the introduction of ‘FAP resulted in a drop in FFA accumulation of up to 90% (for C18:0), whilst for C16:0 there was only a 60% reduction (Fig. 6C). Despite repeated trials, the recovery in the measurable fatty acid to alkane conversion remained poor for C16:0 in comparison to C18:0 and the other pathways tested in *Synechocystis* 6803. This may be explained by an impact on ‘TesA accumulation in the constructs also carrying the gene coding for ‘FAP. Nevertheless, the reconstituted eukaryotic algae alkane pathway was more responsive to introduced modifications in the prokaryotic cyanobacterium than in its native host, though this most likely is explained by challenges associated with the release of FFA in the latter.

Although a substantial amount of both alkanes and alkenes were produced by the engineered strains, their performance likely needs to be improved before any application can be considered. Given that no genetically engineered phototrophic microalgae is currently used for commercial purposes (as far as we are aware), and LCA-studies with non-catalytic systems indicate a low predicted energy return on investment (EROI) (Carneiro et al., 2017), also other challenges with commercial algal biotechnology (e.g. contamination, bioreactor cost, energy consumption, etc) will need to be addressed.

## CONCLUSIONS

The different biosynthetic systems presented in this study varied in terms of cellular context, compartmentation, promoters, operon structures and expression platforms, thus precluding a any direct comparison within and between the two species studied. However, the relative conversion efficiencies and absolute functionalities provide for a valid comparison. As such, it could be seen that the conversion of free fatty acids into alkenes by UndB and alkanes by FAP were effective (>50% conversion, for individual fatty acids up to >90% conversion), and that the native FAP pathway in *Chlamydomonas* was amenable to manipulation but that the inability to increase the FFA pool hindered further progress. Consequently, for alkanes, the reconstruction of the eukaryotic algae pathway in the prokaryotic cyanobacteria host provided a more productive system than the partially synthetic pathways in either of the prokaryotic (CAR-ADO) or eukaryotic hosts (TesA-FAP).

This work describes several approaches to employ synthetic metabolism and substantially exceed native capabilities for hydrocarbon biosynthesis in well-establised model cyanobacteria and algae. Although even greater yields have been reported in oleaginous algae and cyanobacteria that are natively endowed to accumulate lipids, the ability to introduce synthetic metabolic pathways in model strains opens up possibilities for tailored choice of both products and hosts. Importantly, the present work is based on first generation strains and further improvement is likely with systematic optimization of both strains and cultivation conditions, including the use of superior engineered or natural enzyme variants.

## ACKNOWLEDGEMENTS

This project has received funding from the European Union’s Horizon 2020 research and innovation programme project PHOTOFUEL under grant agreement No 640720. IY received a PhD scholarship from Indonesia Endowment Fund for Education (LPDP). The authors would also like to thank Dr. Daniel Jaeger for assistance with lipid extraction from *C. reinhardtii*.

## REFERENCES

Ajjawi, I., Verruto, J., Aqui, M., Soriaga, L. B., Coppersmith, J., Kwok, K., Peach, L., Orchard, E., Kalb, R., Xu, W., Carlson, T. J., Francis, K., Konigsfeld, K., Bartalis, J., Schultz, A., Lambert, W., Schwartz, A. S., Brown, R., Moellering, E. R., 2017. Lipid production in Nannochloropsis gaditana is doubled by decreasing expression of a single transcriptional regulator. Nat Biotechnol. 35, 647–652.

Akhtar, M. K., Dandapani, H., Thiel, K., Jones, P. R., 2015. Microbial production of 1-octanol-a naturally excreted biofuel with diesel-like properties. Metabolic Engineering Communications. 2, 1–5.

Akhtar, M. K., Turner, N. J., Jones, P. R., 2013. Carboxylic acid reductase is a versatile enzyme for the conversion of fatty acids into fuels and chemical commodities. Proc Natl Acad Sci U S A. 110, 87–92.

Angermayr, S. A., van der Woude, A. D., Correddu, D., Vreugdenhil, A., Verrone, V., Hellingwerf, K. J., 2014. Exploring metabolic engineering design principles for the photosynthetic production of lactic acid by Synechocystis sp. PCC6803. Biotechnol Biofuels. 7, 99.

Baier, T., Wichmann, J., Kruse, O., Lauersen, K. J., 2018. Intron-containing algal transgenes mediate efficient recombinant gene expression in the green microalga Chlamydomonas reinhardtii. Nucleic Acids Research. gky532–gky532.

Bernard, A., Domergue, F., Pascal, S., Jetter, R., Renne, C., Faure, J. D., Haslam, R. P., Napier, J. A., Lessire, R., Joubès, J., 2012. Reconstitution of Plant Alkane Biosynthesis in Yeast Demonstrates That Arabidopsis ECERIFERUM1 and ECERIFERUM3 Are Core Components of a Very-Long-Chain Alkane Synthesis Complex. Plant Cell. 24, 3106–18.

Carneiro, M. L. N. M., Pradelle, F., Braga, S. L., Gomes, M. S. P., Martins, A. R. F. A., Turkovics, F., Pradelle, R. N. C., 2017. Potential of biofuels from algae: Comparison with fossil fuels, ethanol and biodiesel in Europe and Brazil through life cycle assessment (LCA). Renewable and Sustainable Energy Reviews. 73, 632–653.

Cho, H., Cronan, J. E., 1995. Defective export of a periplasmic enzyme disrupts regulation of fatty acid synthesis. J Biol Chem. 270, 4216–9.

Cook, C., Dayananda, C., Tennant Richard, K., Love, J., 2017. Third-Generation Biofuels from the Microalga, Botryococcus braunii. Biofuels and Bioenergy.

Delrue, F., Li-Beisson, Y., Setier, P. A., Sahut, C., Roubaud, A., Froment, A. K., Peltier, G., 2013. Comparison of various microalgae liquid biofuel production pathways based on energetic, economic and environmental criteria. Bioresour Technol. 136C, 205–212.

Elhai, J., Wolk, C. P., 1988. Conjugal transfer of DNA to cyanobacteria. Methods Enzymol. 167, 747–54.

Eroglu, E., Melis, A., 2010. Extracellular terpenoid hydrocarbon extraction and quantitation from the green microalgae Botryococcus braunii var. Showa. Bioresource Technology. 101, 2359–2366.

Eser, B. E., Das, D., Han, J., Jones, P. R., Marsh, E. N., 2011. Oxygen-independent alkane formation by non-heme iron-dependent cyanobacterial aldehyde decarbonylase: investigation of kinetics and requirement for an external electron donor. Biochemistry. 50, 10743–50.

Gorman, D. S., Levine, R. P., 1965. Cytochrome f and plastocyanin: their sequence in the photosynthetic electron transport chain of Chlamydomonas reinhardi. Proc Natl Acad Sci U S A. 54, 1665–9.

Hu, P., Borglin, S., Kamennaya, N. A., Chen, L., Park, H., Mahoney, L., Kijac, A., Shan, G., Chavarría, K. L., Zhang, C., Quinn, N. W. T., Wemmer, D., Holman, H.-Y., Jansson, C., 2013. Metabolic phenotyping of the cyanobacterium Synechocystis 6803 engineered for production of alkanes and free fatty acids. Applied Energy. 102, 850–859.

Jing, F., Cantu, D. C., Tvaruzkova, J., Chipman, J. P., Nikolau, B. J., Yandeau-Nelson, M. D., Reilly, P. J., 2011. Phylogenetic and experimental characterization of an acyl-ACP thioesterase family reveals significant diversity in enzymatic specificity and activity. BMC Biochem. 12, 44.

Kaczmarzyk, D., Cengic, I., Yao, L., Hudson, E.P., 2018. Diversion of the long-chain acyl- ACP pool in Synechocystis to fatty alcohols through CRISPRi repression of the es- sential phosphate acyltransferase PlsX. Metab. Eng. 45, 59–66.

Kageyama, H., Waditee-Sirisattha, R., Sirisattha, S., Tanaka, Y., Mahakhant, A., Takabe, T., 2015. Improved Alkane Production in Nitrogen-Fixing and Halotolerant Cyanobacteria via Abiotic Stresses and Genetic Manipulation of Alkane Synthetic Genes. Curr Microbiol. 71, 115–20.

Kallio, P., Pasztor, A., Thiel, K., Akhtar, M. K., Jones, P. R., 2014. An engineered pathway for the biosynthesis of renewable propane. Nature Communications. 5:4731.

Kato, A., Takatani, N., Ikeda, K., Maeda, S.I., Omata, T., 2017. Removal of the product from the culture medium strongly enhances free fatty acid production by genetically engineered. Biotechnol. Biofuels 10, 141.

Khara, B., et al., 2013. Production of propane and other short-chain alkanes by structure-based engineering of ligand specificity in aldehyde-deformylating oxygenase. ChemBioChem 14, 1204–1208.

Kindle, K. L., 1990. High-frequency nuclear transformation of Chlamydomonas reinhardtii. Proc Natl Acad Sci U S A. 87, 1228–32.

Lauersen, K. J., Baier, T., Wichmann, J., Wördenweber, R., Mussgnug, J. H., Hübner, W., Huser, T., Kruse, O., 2016. Efficient phototrophic production of a high-value sesquiterpenoid from the eukaryotic microalga Chlamydomonas reinhardtii. Metab Eng. 38, 331–343.

Lauersen, K. J., Kruse, O., Mussgnug, J. H., 2015. Targeted expression of nuclear transgenes in Chlamydomonas reinhardtii with a versatile, modular vector toolkit. Appl Microbiol Biotechnol. 99, 3491–503.

Lauersen, K. J., Wichmann, J., Baier, T., Kampranis, S. C., Pateraki, I., Møller, B. L., Kruse, O., 2018. Phototrophic production of heterologous diterpenoids and a hydroxyfunctionalized derivative from Chlamydomonas reinhardtii. Metabolic Engineering.

Lea-Smith, D. J., Biller, S. J., Davey, M. P., Cotton, C. A., Perez Sepulveda, B. M., Turchyn, A. V., Scanlan, D. J., Smith, A. G., Chisholm, S. W., Howe, C. J., 2015. Contribution of cyanobacterial alkane production to the ocean hydrocarbon cycle. Proc Natl Acad Sci U S A. 112, 13591–6.

Lea-Smith, D. J., Ortiz-Suarez, M. L., Lenn, T., Nürnberg, D. J., Baers, L. L., Davey, M. P., Parolini, L., Huber, R. G., Cotton, C. A., Mastroianni, G., Bombelli, P., Ungerer, P., Stevens, T. J., Smith, A. G., Bond, P. J., Mullineaux, C. W., Howe, C. J., 2016. Hydrocarbons Are Essential for Optimal Cell Size, Division, and Growth of Cyanobacteria. Plant Physiol. 172, 1928–1940.

Lin, F., Das, D., Lin, X. N., Marsh, E. N., 2013. Aldehyde-forming fatty acyl-CoA reductase from cyanobacteria: expression, purification and characterization of the recombinant enzyme. FEBS J. 280, 4773–81.

Liu, X., Sheng, J., Curtiss, R., 2011. Fatty acid production in genetically modified cya-nobacteria. Proc. Natl. Acad. Sci. USA 108, 6899–6904.

Liu, Y., Wang, C., Yan, J., Zhang, W., Guan, W., Lu, X., Li, S., 2014. Hydrogen peroxide-independent production of α-alkenes by OleTJE P450 fatty acid decarboxylase. Biotechnol Biofuels. 7, 28.

Markley, A.L., Begemann, M.B., Clarke, R.E., Gordon, G.C., Pfleger, B.F, 2015. A syn-thetic biology toolbox for controlling gene expression in the cyanobacterium Synechococcus sp. PCC 7002. ACS Synth. Biol. 4, 595–603.

Metzger, P., Largeau, C., 2005. Botryococcus braunii: a rich source for hydrocarbons and related ether lipids. Appl Microbiol Biotechnol. 66, 486–96.

Neupert, J., Karcher, D., Bock, R., 2009. Generation of Chlamydomonas strains that efficiently express nuclear transgenes. Plant J. 57, 1140–50.

Peca, L., Kós, P. B., Máté, Z., Farsang, A., Vass, I., 2008. Construction of bioluminescent cyanobacterial reporter strains for detection of nickel, cobalt and zinc. FEMS Microbiol Lett. 289, 258–64.

Peramuna, A., Morton, R., Summers, M. L., 2015. Enhancing alkane production in cyanobacterial lipid droplets: a model platform for industrially relevant compound production. Life (Basel). 5, 1111–26.

Qiu, Y., Tittiger, C., Wicker-Thomas, C., Le Goff, G., Young, S., Wajnberg, E., Fricaux, T., Taquet, N., Blomquist, G. J., Feyereisen, R., 2012. An insect-specific P450 oxidative decarbonylase for cuticular hydrocarbon biosynthesis. Proc Natl Acad Sci U S A.

Quinn, J. C., Davis, R., 2015. The potentials and challenges of algae based biofuels: a review of the techno-economic, life cycle, and resource assessment modeling. Bioresour Technol. 184, 444–452.

Rasala, B. A., Barrera, D. J., Ng, J., Plucinak, T. M., Rosenberg, J. N., Weeks, D. P., Oyler, G. A., Peterson, T. C., Haerizadeh, F., Mayfield, S. P., 2013. Expanding the spectral palette of fluorescent proteins for the green microalga Chlamydomonas reinhardtii. Plant J. 74, 545–56.

Rude, M. A., Baron, T. S., Brubaker, S., Alibhai, M., Del Cardayre, S. B., Schirmer, A., 2011. Terminal olefin (1-alkene) biosynthesis by a novel p450 fatty acid decarboxylase from Jeotgalicoccus species. Appl Environ Microbiol. 77, 1718–27.

Ruffing, A.M., 2014. Improved free fatty acid production in cyanobacteria with Synechococcus sp. PCC 7002 as host. Front. Bioeng. Biotechnol. 2, 17.

Rui, Z., Harris, N. C., Zhu, X., Huang, W., Zhang, W., 2015. Discovery of a Family of Desaturase-Like Enzymes for 1-Alkene Biosynthesis. ACS Catalysis. 5, 7091–7094.

Rui, Z., Li, X., Zhu, X., Liu, J., Domigan, B., Barr, I., Cate, J. H., Zhang, W., 2014. Microbial biosynthesis of medium-chain 1-alkenes by a nonheme iron oxidase. Proc Natl Acad Sci U S A. 111, 18237–42.

Schirmer, A., Rude, M., Li, X., Popova, E., del Cardayre, S., 2010. Microbial biosynthesis of alkanes. Science. 329, 559–62.

Sheppard, M. J., Kunjapur, A. M., Prather, K. L. J., 2016. Modular and selective biosynthesis of gasoline-range alkanes. Metab Eng. 33, 28–40.

Sorigué, D., Légeret, B., Cuiné, S., Blangy, S., Moulin, S., Billon, E., Richaud, P., Brugière, S., Couté, Y., Nurizzo, D., Müller, P., Brettel, K., Pignol, D., Arnoux, P., Li-Beisson, Y., Peltier, G., Beisson, F., 2017. An algal photoenzyme converts fatty acids to hydrocarbons. Science. 357, 903–907.

Storch, M., Casini, A., Mackrow, B., Fleming, T., Trewhitt, H., Ellis, T., Baldwin, G. S., 2015. BASIC: A New Biopart Assembly Standard for Idempotent Cloning Provides Accurate, Single-Tier DNA Assembly for Synthetic Biology. ACS Synth Biol. 4, 781–7.

Wang, W., Liu, X., Lu, X., 2013. Engineering cyanobacteria to improve photosynthetic production of alka(e)nes. Biotechnol Biofuels. 6, 69.

Warui, D. M., Pandelia, M. E., Rajakovich, L. J., Krebs, C., Bollinger, J. M., Booker, S. J., 2015. Efficient delivery of long-chain fatty aldehydes from the Nostoc punctiforme acyl-acyl carrier protein reductase to its cognate aldehyde-deformylating oxygenase. Biochemistry. 54, 1006–15.

Wichmann, J., Baier, T., Wentnagel, E., Lauersen, K. J., Kruse, O., 2018. Tailored carbon partitioning for phototrophic production of (E)-α-bisabolene from the green microalga Chlamydomonas reinhardtii. Metab Eng. 45, 211–222.

Work, V.H., Melnicki, M.R., Hill, E.A., Davies, F.K., Kucek, L.A., Beliaev, A.S., et al., 2015. Lauric acid production in a glycogen-less strain of Synechococcus sp. PCC 7002. Front Bioeng. Biotechnol. 3.

Yunus, I.Y., Jones, P.R., 2018. Photosynthesis-dependent biosynthesis of medium chain-length fatty acids and alcohols. Metab Eng. 49, 59–68.

Zhou, Y. J., Hu, Y., Zhu, Z., Siewers, V., Nielsen, J., 2018. Engineering 1-Alkene Biosynthesis and Secretion by Dynamic Regulation in Yeast. ACS Synth Biol. 7, 584–590.

Zhu, Z., Zhou, Y. J., Kang, M. K., Krivoruchko, A., Buijs, N. A., Nielsen, J., 2017. Enabling the synthesis of medium chain alkanes and 1-alkenes in yeast. Metab Eng. 44, 81–88.

